# Freeze-frame imaging of dendritic calcium signals with TubuTag

**DOI:** 10.1101/2020.11.30.403311

**Authors:** Alberto Perez-Alvarez, Florian Huhn, Céline D. Dürst, Andreas Franzelin, Paul Lamothe-Molina, Thomas G. Oertner

## Abstract

The extensive dendritic arbor of neurons is thought to be actively involved in the processing of information. Dendrites contain a rich diversity of ligand- and voltage-activated ion channels as well as metabotropic receptors. In addition, they are capable of releasing calcium from intracellular stores. Under specific conditions, large neurons produce calcium spikes that are locally restricted to a dendritic section. To investigate calcium signaling in dendrites, we introduce TubuTag, a genetically encoded calcium sensor anchored to the cytoskeleton. TubuTag integrates cytoplasmic calcium signals by irreversible photoconversion from green to red fluorescence when illuminated with violet light. To image the mm-long dendritic tree of pyramidal neurons at subcellular resolution, we used a custom two-photon microscope with a large field of view. As the read-out of fluorescence can be performed several hours after photoconversion, TubuTag will help investigating dendritic signal integration and calcium homeostasis in large populations of neurons.

## Introduction

When a cluster of excitatory synapses is simultaneously activated on a basal dendrite of a pyramidal neuron, the combined depolarization triggers a local NMDA spike (Schiller et al., 2000). In addition to NMDA receptors, voltage-gated calcium channels also contribute to local dendritic depolarization and calcium spikes (Losonczy and Magee, 2006; Remy and Spruston, 2007; Takahashi et al., 2020). These nonlinear processes may endow dendrites with the capability to serve as computational subunits (Polsky et al., 2004) and gate the output of cortical neurons during perception of sensory stimuli (Takahashi et al., 2020). As the dendritic arbor of even a single cortical neuron spans several square millimeters and dendritic calcium spikes are stochastic, capturing these rare events with laser scanning microscopy is technically challenging. Most of our knowledge is based on painstaking patch-clamp recordings from individual dendrites or simulating synaptic activity by local glutamate uncaging (Antic et al., 2010; Major et al., 2013). Optical recording *in vivo* requires *a priori* knowledge of input sites since imaging is typically limited to a single plane containing a few dendrites (Helmchen et al., 1999; Takahashi et al., 2020). To generate a lasting record of synaptic activity that can be read-out later, we previously developed SynTagMA (Perez-Alvarez et al., 2020), a synaptically-targeted variant of CaMPARI-2 (Moeyaert et al., 2018). This calcium integrator photoconverts irreversibly from green to red if it is bound to calcium and simultaneously illuminated by violet light (Fosque et al., 2015; Moeyaert et al., 2018). We reasoned that immobilizing the calcium integrator by targeting it to the dendritic cytoskeleton would allow a similar freezing of dendritic calcium events.

To image large populations of neurons at high resolution, low-magnification high-NA water immersion objectives are available from all major microscope manufacturers. However, it is challenging to integrate them into commercial two-photon microscopes as they require a large diameter laser beam scanned at large angles with minimal wave front error. In addition, the detection pathway has to conserve the etendue of the objective (or condenser, respectively) in order to count all emitted photons that enter the optical system. We describe a custom-built two-photon microscope designed to meet the following requirements: 1) 800 × 800 μm field of view (FOV) with good off-axis resolution; 2) efficient detection of green and red fluorescence, simultaneously through the objective and through the condenser; 3) resonant scanning for fast frame rates; 4) protection of photomultiplier tubes (PMTs) during photoconversion (405 nm light); 5) temperature control to maintain brain tissue at 37°C throughout the experiment.

Stable cytoskeletal anchoring and low turnover of TubuTag are essential features to preserve information about subcellular calcium distributions for later read-out by two-photon microscopy. Using hippocampal slice cultures virally transduced with TubuTag as a test system, we show that electrical stimulation of Schaffer collateral axons induces calcium hot spots in dendrites of CA1 neurons. In fixed tissue, the contrast between active and inactive neurons and dendrites can be further enhanced by an antibody that recognizes only the red form of TubuTag.

## Results

Our new sensor is based on the fluorescent protein mEos (Wiedenmann et al., 2004) that undergoes irreversible photoconversion from green to red when illuminated with violet light (405 nm). Previously, mEos had been fused to a calcium-dependent conformation switch (calmodulin-M13) to create integrating sensors that can only be photoconverted when bound to calcium (CaMPARI, CaMPARI2) (Fosque et al., 2015; Moeyaert et al., 2018). To generate TubuTag, we fused CaMPARI2 in-frame to *α*-tubulin. We also created a high affinity version based on CaMPARI2_F391W_L398V (Moeyaert et al., 2018). Expressed in hippocampal neurons by single cell electroporation, dendrites and somata showed bright green fluorescence. Dendritic spines, which rarely contain microtubules and are stabilized by f-actin instead, were largely invisible (Fig. 1a). As the green fluorescence of the CaMPARI moiety dims upon calcium binding, it can be used as an acute calcium sensor. To characterize the calcium sensitivity of TubuTag, we induced back-propagating action potentials (bAPs) by somatic current injections and simultaneously measured green fluorescence intensity in the apical dendrite by two-photon microscopy (Fig. 1b). The dimming response increased with the number of bAPs in a 100 Hz burst, following a sigmoidal curve in a log dose-response plot (Fig. 1c). We found that our tubulin fusion constructs followed the binding curves of the parent CaMPARI variants (Fig. 1d) up to 10 bAPs, but reached saturation slightly earlier (−62% ΔF/F after 30 bAPs). For this study, we used the lowest affinity TubuTag (with CaMPARI2) to minimize baseline photoconversion in inactive cells. CaMPARI2 is also much brighter and shows more efficient photoconversion (larger dynamic range) than CaMPARI1 (Moeyaert et al., 2018).

**Fig. 1.**
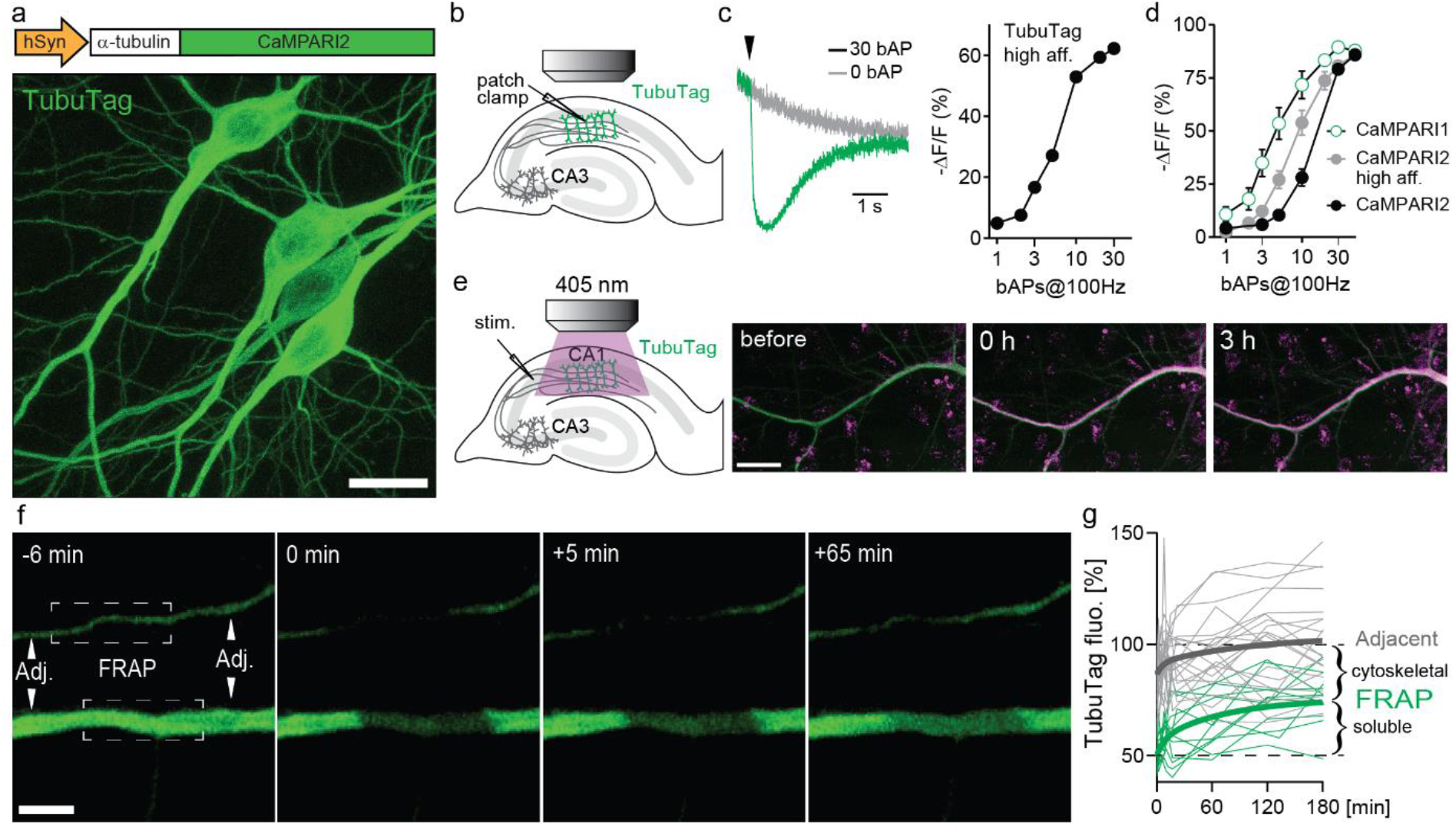
Design and performance of TubuTag. **a,** TubuTag was created by fusing α-tubulin driven by the human synapsin-1 promoter to the calcium integrator CaMPARI2. Expression was tested by single-cell electroporation in CA1. Note absence of TubuTag from dendritic spines that are stabilized by f-actin. Scale bar: 20 μm. **b,** Back-propagating action potentials (bAP) were induced in a TubuTag-expressing neuron while monitoring green fluorescence in the apical dendrite. **c,** Dimming of high affinity TubuTag upon bAP generation through whole-cell patch clamp (2 neurons). Arrowhead indicates time of AP burst. Dimming increases with the number of bAPs**. d**, Dimming of three soluble CaMPARI variants upon bAP induction. **e,** High frequency synaptic stimulation via extracellular electrode (three 0.2 ms pulses, 100 Hz) in *stratum radiatum* followed by a long violet light pulse (405 nm, 14 mW/mm^−2^, 500 ms duration, 1 s delay) led to strong photoconversion of TubuTag (10 pair repeats at 0.033 Hz). Converted TubuTag signal (red) in dendrites remained stable for over 3 hours (2 neurons). Scale bar: 20 μm. **f,** Turnover of TubuTag was assessed by fluorescence recovery after photobleaching (FRAP). Fluorescence at bleached (squares) and adjacent (arrowheads) dendritic areas was measured and followed over time. Scale bar: 5 μm. **g,** Dendritic fluorescence normalized to F_0_ fitted by a double exponential for bleached (τ_fast_ = 8 min; τ_slow_ = 75 min, n = 12 dendrites, 6 neurons) and adjacent areas.

Illumination with long violet light (405 nm) pulses during extracellular high-frequency stimulation of CA3 axons induced strong photoconversion of CA1 neurons expressing TubuTag (Fig. 1e). The red label showed no sign of decay after 3 h, indicating irreversible photoconversion and slow protein turnover. Therefore, we expect red/green ratio differences between active and inactive neurons to be stable for several hours.

To determine the fraction of TubuTag integrated in stable microtubules vs soluble TubuTag monomers, we measured fluorescence recovery after photobleaching (FRAP, Fig. 1f). After 3 hours, ^~^50% of the bleached fluorescence had recovered (Fig. 1g), consistent with the slow turnover of microtubules in neurons (Kahn et al., 2018). Adjacent regions showed a small and transient dip in fluorescence intensity, consistent with diffusional exchange of bleached and unbleached monomers. Thus, if subcellular resolution is the goal of the experiment, the red/green ratio (R/G) should be determined within ~20 min after photoconversion.

Objectives with high numerical aperture (NA) and low magnification are ideal to image an entire neuron at high resolution in a single stack of images. To use the full potential of these objectives for two-photon imaging, we designed a 3-axis scanner (resonant-galvo-galvo) with relay optics based on parabolic mirrors. In our design, the scan lens - tube lens assembly provides 4.3 × magnification and is horizontally oriented, allowing for a compact detection assembly directly on top of the objective. Below the stage, a second optical system detects fluorescence through a 1.4 NA oil immersion condenser and provides IR-Dodt contrast for electrophysiological experiments. During photoconversion, both light detection pathways are protected by electronic shutters with 45 mm free aperture (NS45B, Uniblitz). The oil-immersion condenser is heated to 33°C by a ring of Peltier elements, with the cool side in contact with the PMT assembly. The xy-stage and z-focus are motorized (Sutter MPC200). A magnetic holder allows switching objectives with different working distances for different types of experiments (Olympus XLPlan 25x 1.0 NA, 4 mm WD; Nikon 16x 0.8 NA, 3 mm WD; Olympus XLPlan 10X 0.6 NA, 8 mm WD).

For most experiments, we used a Nikon 16x 0.8 NA objective, which provided a good compromise between FOV (1.7 × 1.7 mm), optical resolution (0.5 × 0.5 × 3 μm^3^) and 45-degree approach angle for patch-clamp or stimulation electrodes. We moved the resonant scan field (875 × 875 μm) within the large FOV using the two galvanometric mirrors.

In order to express our sensor in a population of neurons, we packaged TubuTag into an AAV9 vector and injected it into the CA1 region of hippocampal organotypic slices using a Picospritzer (Fig. 3a). Schaffer collateral axons were stimulated with an extracellular electrode in *stratum radiatum* (paired pulses at 20 Hz), shortly followed by a short violet light pulse delivered through the imaging objective. After repeating this protocol 50 times, many dendrites in the vicinity of the electrode turned red while more distant dendrites remained green (Fig. 3b). Strongly converted dendrites could be found next to non-converted dendrites, suggesting that not all neurons produced dendritic calcium spikes (Fig. 3c). Conversion efficiency was not different in superficial vs deep dendrites, indicating good penetration of the 405 nm photoconversion light (Fig. 3d). The high transduction efficiency of AAV9 allowed us to monitor compartmentalized calcium signals at full density in a large volume fraction of CA1 in a single experiment. As TubuTag expression was driven by the synapsin-1 promoter, expression was not restricted to pyramidal cells but included interneurons. Interestingly, we found that some interneuron dendrites showed ‘hot spots’ of high photoconversion (Fig. 3f). This inhomogeneity, which has also been noted in live calcium imaging experiments (Soler-Llavina and Sabatini, 2006; Francavilla et al., 2019; Topolnik and Camiré, 2019), points to highly compartmentalized calcium signaling.

**Fig. 2:**
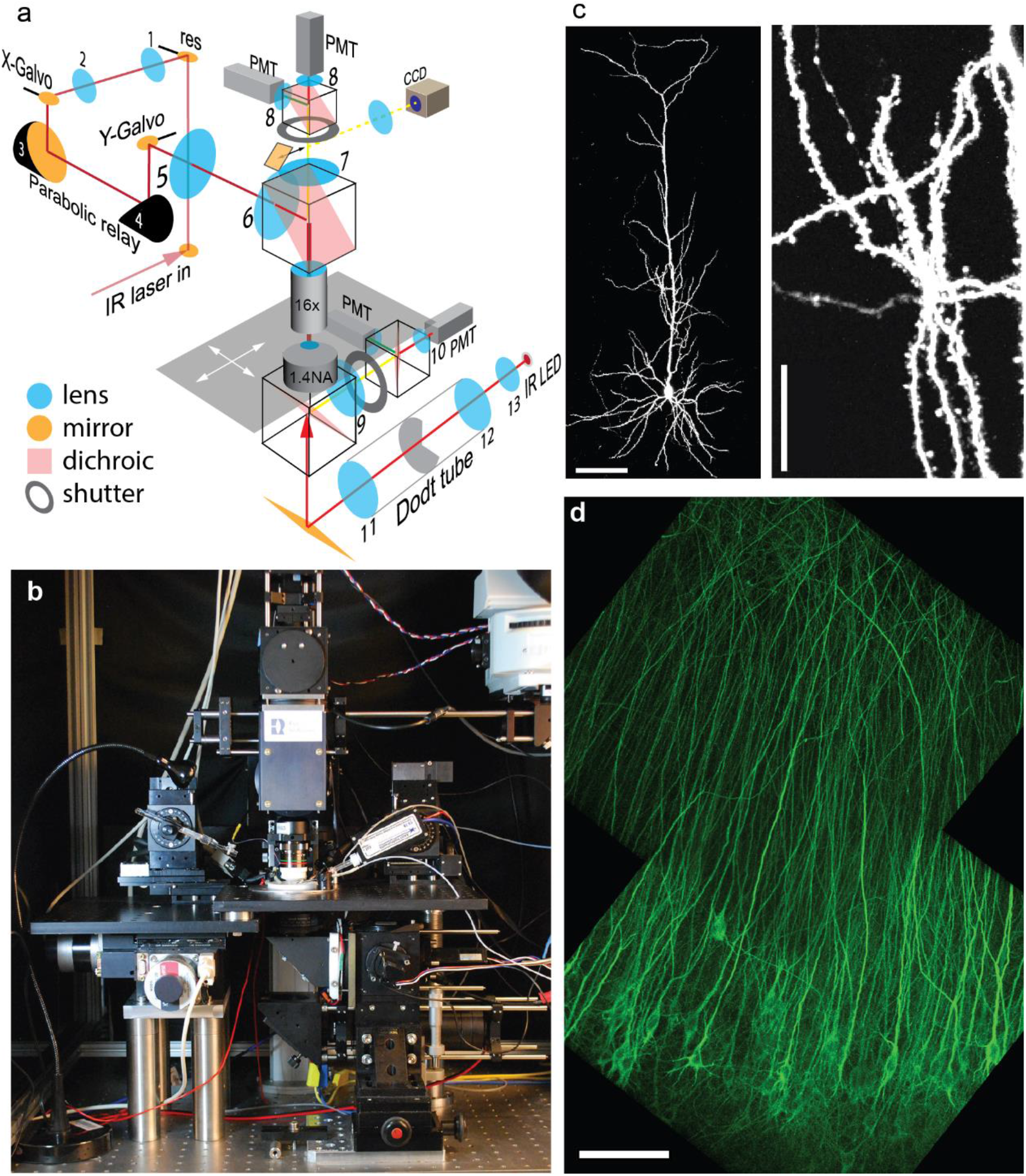
Design and performance of the Rapid3D microscope. **a,** The pulsed IR laser is scanned via resonant (res) and galvanometric scan mirrors (X-Galvo, Y-Galvo) through a scan lens – tube lens assembly (5-6) and a 25 × 1.0 NA objective. The first relay (1-2) provides a magnification factor M = 1.3, the parabolic relay (3-4) has M = 1, the scan lens - tube lens assembly (5-6) provides M = 4.5. Fluorescence is detected by 4 photomultiplier tubes (PMT) through red and green band pass filters. A substage IR LED provides Dodt contrast illumination (CCD camera) for patch clamp experiments. **b,** Frontal view of the Rapid3D microscope. **c,** Image (maximum intensity projection) of a single CA1 pyramidal neuron expressing a cytoplasmic red fluorescent protein (tdimer2). Dendritic spines and axonal boutons are well resolved. Scale bars: 100 μm (left), 20 μm (right). **d,** Virally transfected CA1 pyramidal cells expressing TubuTag. Note apparent absence of dendritic spines as they rarely contain microtubules. Scale bar: 100 μm.

**Fig. 3:**
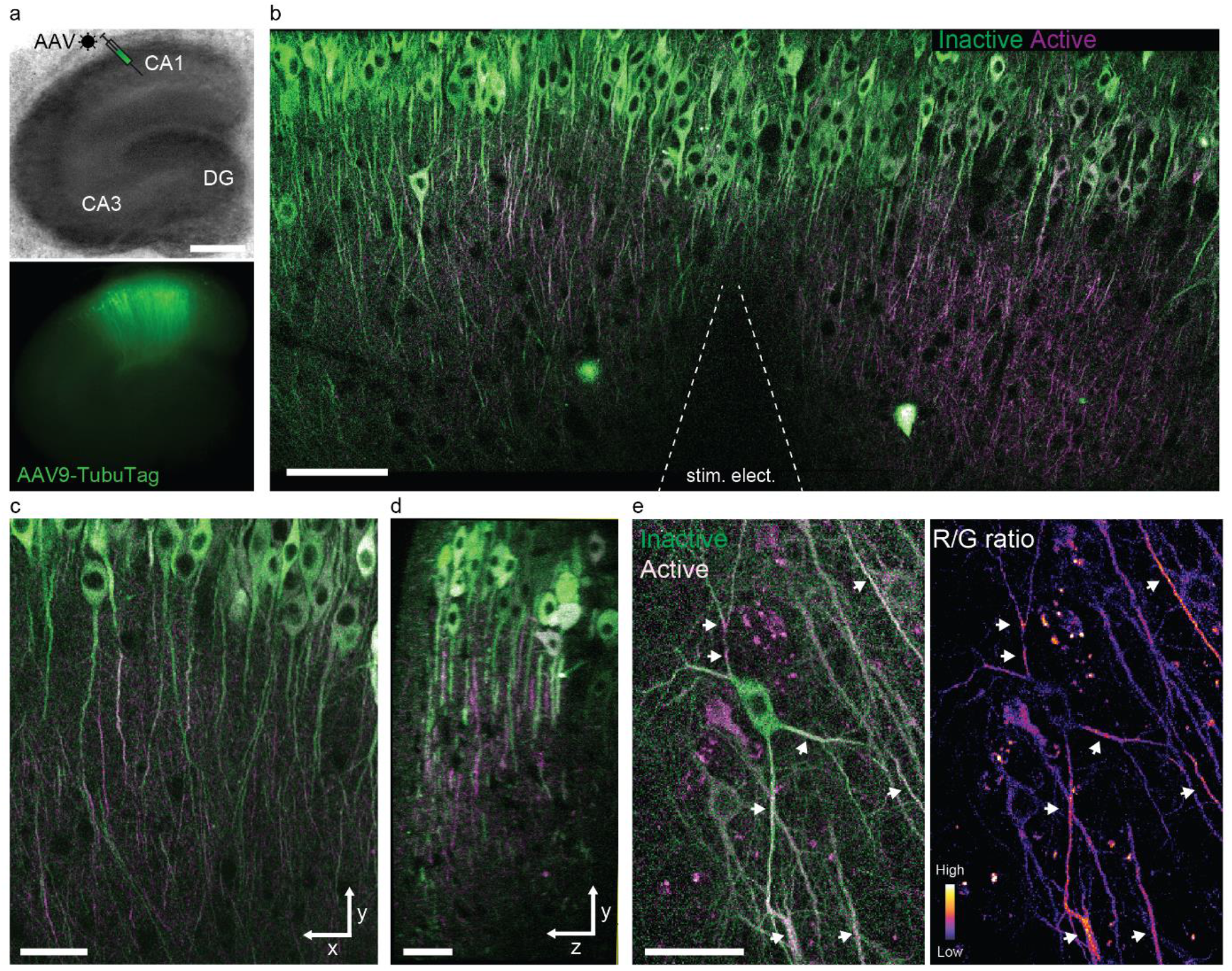
Photoconversion of TubuTag by extracellular stimulation paired with violet light. **a,** Rat hippocampal slice culture microinjected with AAV9-syn-TubuTag in the CA1 region. Scale bar: 500 μm. **b,** Extracellular stimulation in *stratum radiatum* was paired with short violet illumination (405 nm, 11 mW/mm^−2^, 200 ms duration, 500 ms delay, 50 pair repeats at 0.066 Hz) through the objective to induce photoconversion of calcium-bound TubuTag. Scale bar: 100 μm. **c,** Detail of b, showing strong differences in photoconversion between neighboring dendrites. Scale bar: 50 μm. **d,** Side view of volume shown in c. Scale bar: 30 μm. Note that photoconversion is not restricted to superficial dendrites (left), but independent of depth. **e,** Interneuron expressing TubuTag in *stratum radiatum* after photoconversion. Arrowheads denote dendritic sections where strong photoconversion occurred. Scale bar: 50 μm. Low to high converted dendrites are shown by pseudocoloring the red-to-green ratio.

It would be attractive to induce TubuTag conversion in intact animals during behavior and image the frozen calcium signal at high resolution *ex vivo*. Chemical fixation, however, invariably leads to some loss of fluorescence, reducing imaging contrast. For *ex-vivo* detection of active dendrites, it would be advantageous to specifically enhance the red fluorescence by immunostaining. A specific antibody against the red form of CaMPARI has been developed (Moeyaert et al., 2018). As a proof of principle, we tested this antibody on slices where TubuTag neurons had been strongly photoconverted with long violet light pulses while being driven to spike. In the live tissue, we could visualize layer-specific photoconversion across the whole CA1 region (Fig. 4a). After fixation, we used the primary anti-CaMPARI-red antibody and a far-red secondary antibody (AF648) to selectively enhance the photoconverted signal. Anti-CaMPARI-red highlighted exactly the same neurons we previously identified as converted, indicating excellent specificity for the photoconverted form of TubuTag (Fig. 4b, c).

**Fig. 4:**
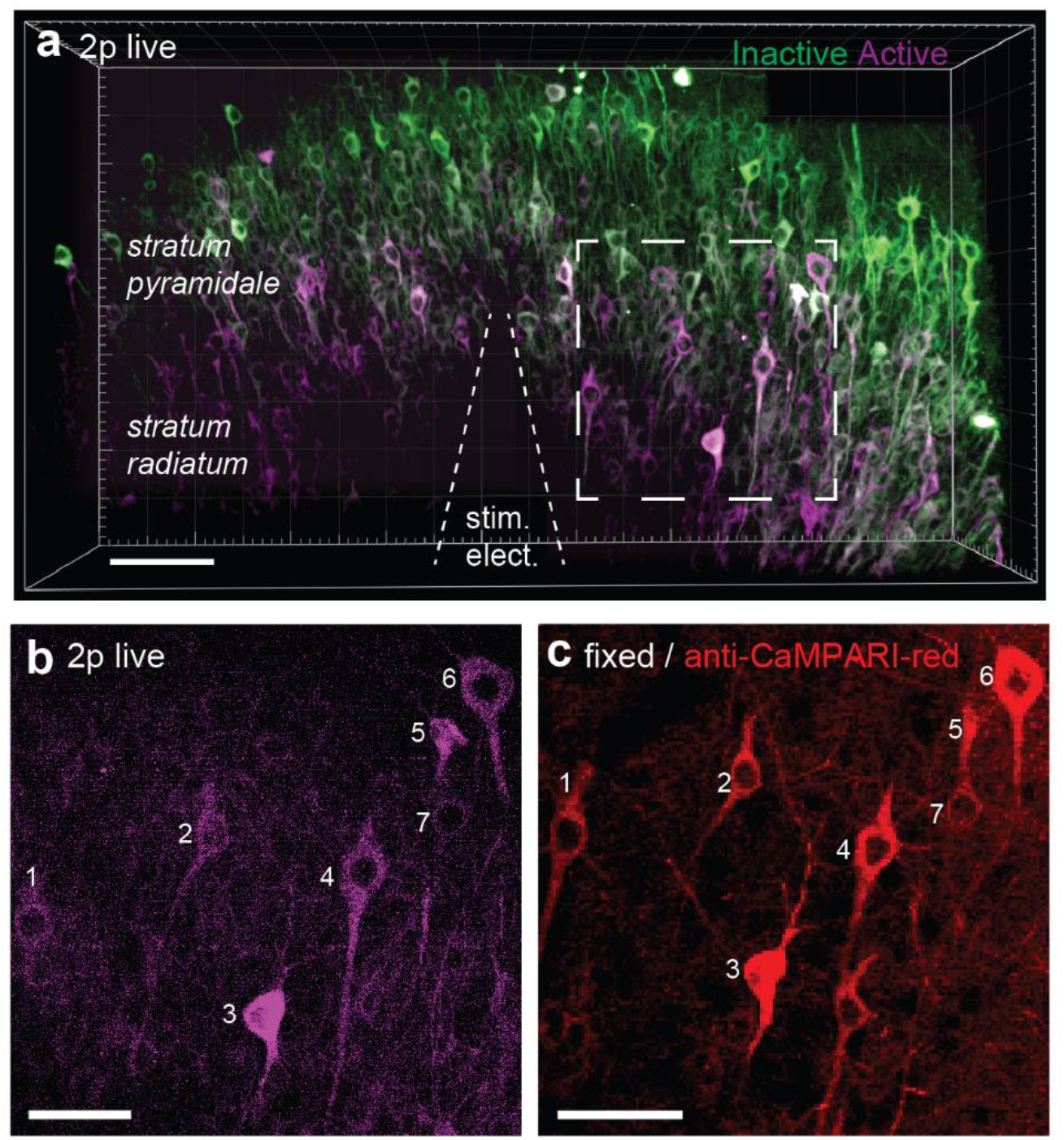
TubuTag *ex vivo* immunodetection. **a,** Three-dimensional display of CA1 area of hippocampal slice virally transfected with TubuTag after photoconversion. Electrical stimulation at 37°C followed by long violet light illumination (405 nm, 11 mW/mm^−2^, 500 ms duration, 1 s delay, 50 pair repeats at 0.1 Hz). Scale bar: 100 μm. **b,** Inset in a (maximum projection) showing photoconverted neurons (magenta). Scale bar: 50 μm. **c,** Confocal image of same region in b after fixation and staining against red CaMPARI (i.e., photoconverted). Scale bar: 50 μm.

## Discussion

Calcium signals provide a unique window into neuronal physiology. From the release of neurotransmitter, induction of synaptic plasticity, complex spike bursts, to immediate early gene expression: all significant events in the life of a neuron are accompanied by calcium transients. Two-photon microscopy is a powerful tool to monitor such intracellular calcium signals at high resolution in intact brain tissue. Since fluorescence is sampled point-by-point, however, imaging of large and densely labeled volumes at the speed necessary to catch physiological calcium signals in subcellular detail is not really possible with this technique.

The calcium integrator CaMPARI irreversibly labels active neurons in a time window defined by violet illumination. This was a revolutionary concept, since it completely dissociated the labeling event from the optical recording of the results. Obviously, the speed of the microscope becomes much less important once the functional signal is frozen in time. Since CaMPARI is free to diffuse inside a neuron, any subcellular gradients in photoconversion rapidly dissipate and the location of active inputs is lost. By fusing CaMPARI2 to tubulin, we immobilized the calcium reporter, providing a lasting record of high [Ca^2+^] areas across the dendritic arbor. TubuTag imaging is relatively straightforward, as a single wavelength (1040 nm) can be used to excite both green and red species simultaneously. Consequently, there is no need to register green and red images as both are excited by a single laser focus.

If a TubuTag-expressing neuron spikes at high frequency during violet light application, not only input sites, but also other parts of the dendrite could become photoconverted. For precise localization of active inputs, it may thus be useful to co-express TubuTag with a light-gated potassium or chloride channel (Wietek et al., 2015; Beck et al., 2018) to prevent action potential generation during the photoconversion period.

A topic of considerable interest is the possibility that synapses with synchronous activity are able to stabilize each other when situated close together, leading over time to a clustered input organization (Kleindienst et al., 2011; Iacaruso et al., 2017). In principle, an activity tag exclusively located at excitatory synapses like postSynTagMA (Perez-Alvarez et al., 2020) could be used to investigate the input organization of individual neurons, but photoconversion of a group of spines could also result from a local dendritic calcium event. Combining SynTagMA and TubuTag might be a way to resolve this ambivalence: red spines on a green section of dendrite would be incontrovertible evidence for synchronous synaptic activity.

The most important feature of TubuTag is the hour-long preservation of subcellular [Ca^2+^] gradients in live tissue. We envision that a head-mounted violet LED could be used to trigger photoconversion in intact animals, a situation where high-resolution imaging is very challenging. As we show, the signal-to-noise ratio in fixed tissue can be considerably improved by an antibody that recognizes only the red form of TubuTag. Compared to acute calcium imaging *in vivo*, which usually requires head fixation, TubuTag may provide a way to map cortical activity in freely moving animals.

## Methods

### Plasmid construction

TubuTag is CaMPARI2 (Moeyaert et al., 2018) fused to human α-tubulin, high affinity TubuTag contains 2 point mutations in the CaMPARI2 domain, F391W and L398V. To generate high affinity TubuTag, we replaced mCherry from pcDNA3.1 mCherry-human-alpha-tubulin (a gift from Marina Mikhaylova) with the CaMPARI moiety from preSynTagMA (Perez-Alvarez et al., 2020); ref. Addgene 119738) using HindIII and Bsp 1407I restriction sites. To insert the CaMPARI-tubulin fusion construct into a pAAV backbone, we first replaced in the C-terminus the restriction site XhoI by HindIII using PCR. Subsequently, we used NheI and HindII to insert it into the pAAV backbone of pAAV-syn-ChR2ETTC (Bernd et al., 2011, ref Addgene 101361). To create TubuTag, we mutated residues W391 and V398 by overlap extension PCR to obtain a W391F_V398L variant (CaMPARI2). Briefly, we designed primers to amplify two DNA segments containing both the point mutations, namely between NheI and V398L, W391F and BoxI. The resulting amplified segments with sizes 1304 bp and 728 bp were overlaid, resulting in a 1968 bp segment size that was inserted to replace the segment in TubuTag using restriction sites NheI and BoxI.

### Organotypic hippocampal slice cultures

Hippocampal slice cultures from Wistar rats of either sex were prepared at postnatal day 4–7 as described (Gee et al., 2017). Briefly, rats were anesthetized with 80% CO_2_ 20% O_2_ and decapitated. Hippocampi were dissected in cold slice culture dissection medium containing (in mM): 248 sucrose, 26 NaHCO_3_, 10 glucose, 4 KCl, 5 MgCl_2_, 1 CaCl_2_, 2 kynurenic acid and 0.001% phenol red. pH was 7.4, osmolarity 310-320 mOsm kg^−1^, and solution was saturated with 95% O_2_, 5% CO_2_. Tissue was cut into 400 μM thick sections on a tissue chopper and cultured at the medium/air interface on membranes (Millipore PICMORG50) at 37° C in 5% CO_2_. No antibiotics were added to the slice culture medium which was partially exchanged (60-70%) twice per week and contained (for 500 ml): 394 ml Minimal Essential Medium (Sigma), 100 ml heat inactivated donor horse serum (Sigma), 1 mM L-glutamine (Gibco), 0.01 mg ml^−1^ insulin (Sigma), 1.45 ml 5 M NaCl (Sigma), 2 mM MgSO_4_ (Fluka), 1.44 mM CaCl_2_ (Fluka), 0.00125% ascorbic acid (Fluka), 13 mM D-glucose (Fluka). Wistar rats were housed and bred at the University Medical Center Hamburg-Eppendorf. All procedures were performed in compliance with German law and according to the guidelines of Directive 2010/63/EU. Protocols were approved by the Behörde für Justiz und Verbraucherschutz (BJV) - Lebensmittelsicherheit und Veterinärwesen, of the City of Hamburg.

### Neuronl transduction

CA1 neurons in rat organotypic hippocampal slice culture (DIV 17-21) were transfected by single-cell electroporation (Wiegert, Gee and Oertner, 2017). Thin-walled pipettes (~10 MΩ) were filled with intracellular K-gluconate based solution into which plasmid DNA was diluted to 20 ng μl^−1^. Pipettes were positioned against neurons and DNA was ejected using an Axoporator 800A (Molecular Devices) with 50 hyperpolarizing pulses (−12 V, 0.5 ms) at 50 Hz. Experiments were conducted 3-5 days after electroporation.

For viral transduction of hippocampal slices, AAV2/9 viral vectors containing TubuTag variants under the control of the synapsin promoter was prepared at the UKE vector facility. Working in a laminar air flow hood, organotypic hippocampal slices were microinjected at DIV 7-11 (Wiegert et al., 2017b). A pulled glass pipette was backfilled with 1 μl of the viral vector and the tip inserted in the hippocampal CA1 area. A picospritzer (Science Products) coupled to the pipette was used to deliver 3-4 short (50 ms) low pressure puffs of viral vector into the tissue. Injected slices were taken back to the incubator and imaged in the multiphoton microscope 15-21 days later.

### Two-photon imaging and photoconversion

For fast two-photon imaging, we used a customized version of the Rapid3Dscope (Rapp OptoElectronic GmbH), a large field of view two-photon microscope equipped with relayed resonant-galvo-galvo scanners (RGG). The microscope is controlled by the open access software ScanImage 2017b (Vidrio; Pologruto et al., 2003). To simultaneously excite both the green and red species of TubuTag, we employed a pulsed Ti:Sapphire laser (Chameleon Ultra II, Coherent) tuned to 1040 nm.

Red and green fluorescence was detected through upper and lower detection path with one pair of photomultiplier tubes (PMTs, H7422P-40SEL, Hamamatsu) on each side. The lower detection has an oil immersion condenser (1.4 NA, Olympus), a 560 DXCR dichroic mirror, and 525/50 (green) and 607/70 (red) emission filters (Chroma Technology). The upper detection has a removable handle and adaptor to accommodate several magnification objectives (CFI75 LWD, 16x, 0.8 NA, Nikon), the main short-pass dichroic (Chroma ZT775sp-2p) and a dichroic mirror (Chroma T565lpxr) with red and green band path emission filters (Chroma ET525/70m-2p, Chroma ET605/70m-2p). Excitation light was blocked by short-pass filters (Chroma ET700SP-2P).

For extracellular synaptic stimulation, a glass monopolar electrode filled with extracellular solution was placed in *stratum radiatum*. Paired 0.2 ms pulses (40 ms inter stimulus interval) were delivered using an ISO-Flex stimulator (A.M.P.I.) at 0.1 Hz. Overview images were acquired, covering an area close to the electrode tip. The galvo - resonant scanner relay allowed z-stack acquisition covering a brain volume of 435 × 435 × 100 μm^3^ in 2 min at 1024 pixel resolution and 1 μm z-step. Repeated triggering of electrical stimulation and violet light illumination (405 nm, CoolLED pE4000) through the epifluorescence path and objective was done by the electrophysiology software Wavesurfer (Janelia Research). The resonant scanner was engaged before image acquisition start, and laser power was controlled by a Pockels cell (Conoptics). Electronic shutters (Uniblitz) were in place both at the beam path to protect the user and the sample, and in the detection paths to protect PMTs between acquisition periods. Series of 2-3 images acquired to cover a mm-long brain area were registered using the pairwise stitching plugin from ImageJ (Preibisch et al., 2009).

For suprathreshold stimulation and photoconversion, three 0.2 ms pulses at 100Hz were applied with a glass monopolar electrode followed 1 s later by 500 ms violet light (14mW/mm^−2^). This pairing was repeated ten times at 0.033 Hz.

For calcium sensitivity calibration, CA1 neurons expressing TubuTag or CaMPARI variants or 3-5 days after electroporation were patched and brought into whole-cell configuration. Back-propagating action potentials (bAPs) were induced by injecting 1 nA pulses of 1 ms duration in current clamp. Fluorescence in the main apical dendrite was acquired at 500 Hz and changes from basal values were reported as ΔF/F (F-F_0_/F_0_). To depict photoconversion, red and green channels were merged. Alternatively, a ratio image dividing the red by the green channel was obtained with the ratio plus plugin of ImageJ.

### Fluorescence recovery after photobleaching (FRAP)

To estimate the diffusion of TubuTag molecules, we took z-stacks of dendrites in *stratum radiatum* from neurons electroporated with TubuTag and we bleached central parts of dendrites, subsequently following the recovery of fluorescence over time. After obtaining a baseline of 1-2 images, we zoomed (4X) onto the dendrite and performed 10 z-stacks at twice the laser power to achieve ~50% bleaching without signs of tissue damage. Images were subsequently taken at the original settings and magnification at different intervals for 180 min. For analysis of each dendrite, we used a macro written in Fiji (Schindelin et al., 2012) to quantify fluorescence changes over time (Moeyaert et al., 2018) in the bleached area and two immediately adjacent non-bleached areas. Kinetics of recovery after bleaching was fit using nonlinear regression in Prism 8 (GraphPad).

### Immunohystochemistry

After imaging and photoconversion in the 2p microscope, slices were fixed in a PBS solution containing 4% PFA for 30 min at room temperature (RT), washed and kept overnight in PBS at 4°C. Slices were then incubated in blocking buffer (1x PBS, 0.3% TritonX, 5% goat serum) for 2 hours at RT. Next, sections were placed in the primary antibody (1:1000, mouse anti campari2 red 4F6-1 clone, a gift from Eric Schreiter) carrier solution (PBS, 0.3% TritonX, 1% goat serum, 1% BSA) and incubated overnight at 4°C. After 3x washing with 1xPBS for 5 min, the sections were incubated for 2 h at RT in the secondary antibody (1:1000, goat anti mouse Alexa Fluor 647, Life Technologies) carrier solution (1x PBS, 0,3% TritonX, 5% goat serum). Sections were washed 3 × 10 min in 1x PBS, mounted on coverslips using Immu-Mount (Shandon), and imaged with a confocal microscope (Olympus FV 1000) using a 20x oil immersion objective (UPLSAPO 20X NA 0.85). Two-channel images were obtained at 1024×1024 pixel resolution. Excitation/emission spectra and filters were selected using the automatic dye selection function of the Olympus fluoview software (Alexa 488 and Alexa 647).

## Acknowledgements

We thank Iris Ohmert, Sabine Graf and Jan Schröder for excellent technical assistance. AAV vectors were produced by Ingke Braren of the UKE Vector Facility. Anti-red CaMPARI was a gift from Eric Schreiter, mCherry-α-tubulin was a gift from Marina Mikhaylova. The project was supported by the German Federal Ministry of Education and Research (BMBF) through the grant “Rapid3D” (FKZ: 13N14537) in the initiative “KMU-innovativ: Photonik und Quantentechnologien”, and by the German Research Foundation (DFG) through grants SPP 1665 220176618, SFB 936 178316478 and FOR 2419 278170285.

